# Live-cell 3D-SIM of Rift Valley fever virus NSs filaments reveals a polygonal web architecture

**DOI:** 10.1101/2025.07.30.667610

**Authors:** James Dunlop, Peter A. Thomason, Leo M. Carlin, Benjamin G. Davis, Stephen D. Carter

## Abstract

A defining feature of Rift Valley fever virus (RVFV) is the incorporation of the NSs protein into large filamentous assemblies inside infected nuclei ^1^, as judged from fixed specimens. To gain insight into the three-dimensional (3D) structure of NSs filaments within nuclei of live-cells we used genetic-code expansion (GCE) to introduce an unnatural amino acid (uAA) fluorophore into the protein, and coupled this with live-cell structured illumination microscopy (SIM). Our super-resolved images revealed a micron-scale polygonal web of NSs fibres with discrete domain characteristics. Parallel experiments on fixed RVFV-infected cells confirmed that virally-encoded native NSs filaments also display this morphology. Overall, our 3D-SIM analysis sheds new light on the complex large-scale architecture of NSs filaments and provides context for programmable filamentous E3 ligases that promote virus replication.

**Significance:** This work demonstrates how advanced labeling and imaging techniques can reveal unexpected organizational principles in viral protein assemblies. We bio-orthogonally labeled the Rift Valley fever virus NSs protein with bright tetrazine dyes using genetic-code expansion (GCE), that enabled imaging of NSs filaments in live-cells using 3D-SIM. GCE offers key advantages for studying small, genetically fragile viral proteins like NSs. It eliminates the need for bulky fluorescent proteins that can disrupt protein-protein interfaces critical for complex assembly and avoids fixation artifacts associated with traditional immunofluorescence microscopy. This integrative GCE and 3D-SIM approach revealed the complete native architecture of NSs filaments in 3D, dramatically enhancing our ability to visualize intricate polygonal web structures beyond simple linear filaments. The discrete structural domains identified here raise intriguing questions about potential mechanisms for spatially controlling E3 ligase activity in a programmable manner.

## Introduction

Rift Valley fever virus (RVFV) outbreaks cause severe human disease across North Africa and the Arabian Peninsula, and are considered an emerging threat among a small group of priority pathogens included in the Blueprint list by the World Health Organization^2^. The pathogenesis of RVFV is driven by the virally encoded non-structural protein (NSs) that forms filamentous structures inside infected nuclei^1^. Recent eTorts identified the assembly of an NSs filament-FBXO3 E3 ligase complex that was shown to degrade TFIIH and RNA-dependent protein kinase (PKR)^3,4^. Consequences of TFIIH and PKR degradation are crucial for RVFV virulence ^5–7^.

Thin-section transmission electron microscopy (TEM) revealed NSs filaments were composed of bundles of loosely packed, roughly parallel 10 nm wide filaments ^8,9^. This filamentous architecture has been supported by X-ray crystallography ^8^, and cryo-electron microscopy (cryo-EM ^4^. The cryo-EM structure revealed right-handed filaments consisting of three types of NSs-NSs interface crucial for filament formation.

3D-SIM can be conducted with either live or fixed samples. Fixation can distort important structural features, and permeabilisation is required for antibodies to access their intracellular targets. To avoid further disrupting NSs architecture with genetically-encoded fluorescent proteins, we instead used genetic-code expansion (GCE) to label NSs with a bright, minimally disruptive uAA-fluorophore. This approach enabled 3D-SIM imaging of intact NSs filaments in live-cells ^10^. This allowed us to elucidate the native micron-scale architecture of NSs filaments. Using artificial intelligence (AI) powered segmentation ^11,12^, we uncovered an intricate web of NSs fibres that show discrete architectures such as “sheet”, “chain”, and “twisting” morphology.

## Results

We used GCE to incorporate an uAA fluorophore TCO*A (trans-Cyclooct-2-en-L-Lysine) into NSs via amber stop-codon suppression ^10,13^. To guide selection of appropriate amber stop-codon sites we utilized the RVFV NSs cryo-EM map (EMDB:37443) **(Figure 1A)**. We avoided NSs-NSs interface residues, and selected 4 positively charged and 11 polar, uncharged surface exposed residues as candidates **(Figure 1A)**. To test amber suppression, we expressed each of the 15 NSs mutants **(from strain ZH548)** in HEK-293T cells in the presence of pyrrolysine (Pyl) tRNA and a *M*.*mazei*-derived tRNACUAPyl synthetase (see methods), with and without TCO*A. Cell extracts were analysed by Western blotting **(Figure 1B)**, indicating that five candidate amber-suppression site mutants **(K108, S119, S126, S127, Q140-TCO*A)** produced abundant, fragment-free protein consistent in size with full-length WT NSs protein in the presence of TCO*A **(Figure 1B)**.

**Figure 1.**
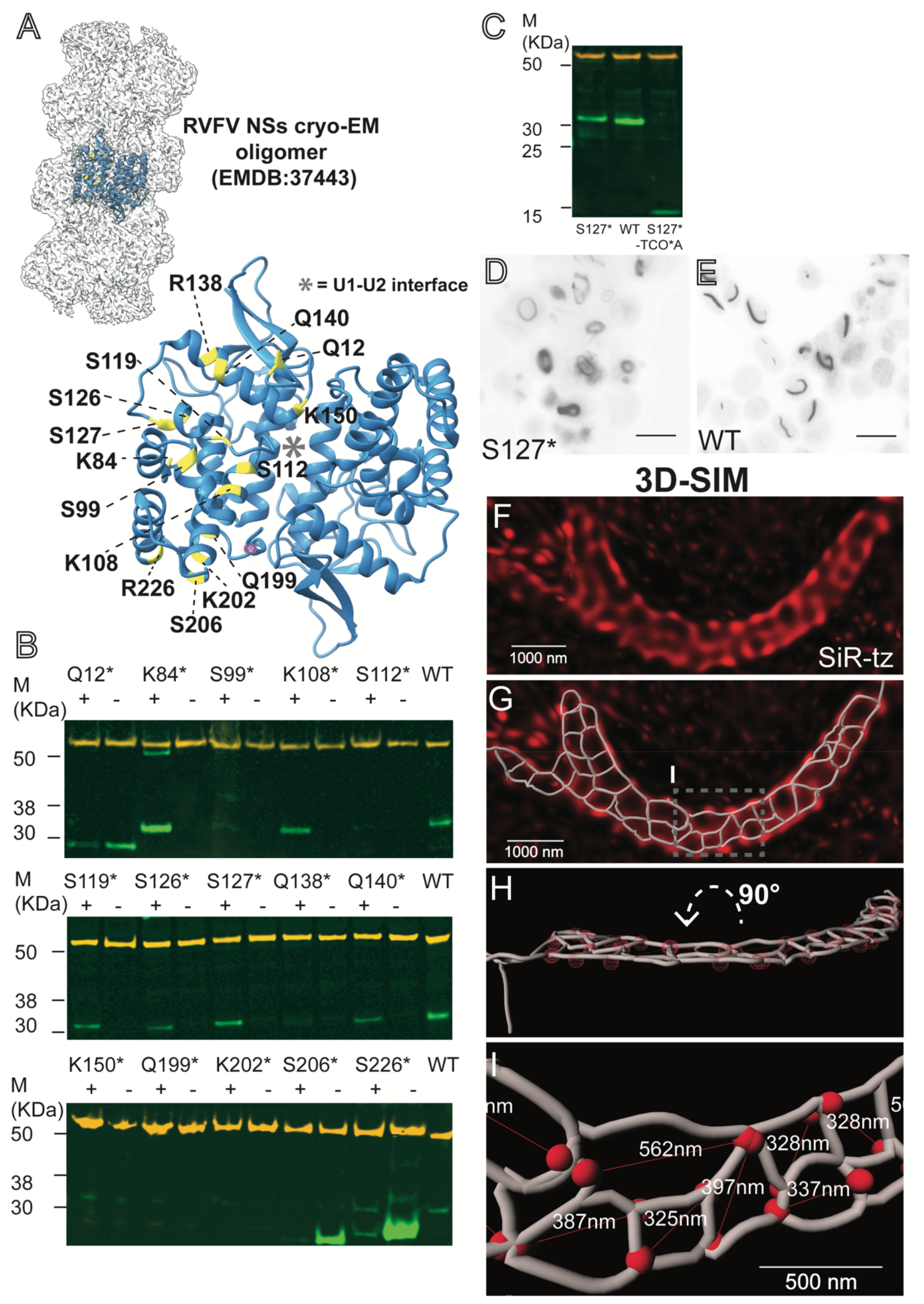
(A) RVFV NSs oligomer^4^. The NSs-NSs dimer is highlighted as a blue ribbon (PDB:8WCM). Stop-codon residues are highlighted in yellow and numbered. (B) Western blot analysis of amber-suppression NSs mutants in HEK-293T cells (denoted as an asterisk). Tubulin in orange. (C) Western blot analysis of amber-suppression in HeLa cells for S127TCO*A (S127*). (D) Wide-field fluorescence image of S127* NSs filaments labeled with CF^®^500 in HEK-293T cells. (E) Immunofluorescence of WT NSs filaments in HEK-293T cells. Scale bars represent 20 µm. (F) Live-cell 3D-SIM analysis of S127* SiR-tz filaments (red) labeled in HeLa cells. (G) A segmentation overlaid on the 3D-SIM image shown in G. (H) The segmentation in H rotated 90 degrees around the X-axis to a more useful perspective. (I) Enlarged view of the gray dashed inset in G with diameters of the polygons measured in the longest dimension.

Next, we independently confirmed protein formation by screening for NSs filament assembly using wide-field fluorescence microscopy. To do this, we used bio-orthogonal click chemistry to label **K108, S119, S126, S127 and Q140-TCO*A** NSs proteins respectively with the CF-500 tetrazine dye. For **Q140TCO*A** and **S127TCO*A**, we observed filaments that were identical in morphology to filaments formed during WT NSs transient overexpression **(Figure 1, compare panels D and E)**. Next, we exploited 3D-SIM and imaged **S127TCO*A** NSs filaments in live-cells. For this we used HeLa cells instead of HEK cells, due to their superior imaging properties, after confirming that full-length **S127TCO*A** NSs protein was expressed **(Figure 1C)**.

Unexpectedly, both live-cell and fixed 3D-SIM imaging of NSs labeled with SiR-tz (red) and Janelia Fluor^®^ 549 (yellow), respectively, revealed large-scale architecture of an open filamentous network. The network contained domains exhibiting intersecting parallel, perpendicular and oblique filaments that produced a series of heterogeneous 3D polygons **(Figure 1F and 2D) (# of filaments imaged in Table 1)**. To further characterise network morphology, we applied an AI powered filament tracer tool in Imaris, with manual validation steps, that delineated an interconnected network of branching fibres and Y-shaped junctions, comprising a polygonal web architecture **(Figure 1F-I)**. This encompassed a variety of morphologies including open lattice structures of flat-sheets **(Figure 1F-I)**, twisting-sheets **(Figure 2F)**, polygon chains **(Figure 2A-C)**, and larger and complex networks **(Figure 2D-F and movie 4)**, some with twisting morphology **(Figure 2G and movie 1 and 2)**.

**Table 1.**

| Filament type | Number of filaments imaged by 3D-SIM |
| --- | --- |
| Live-cell (GCE) | 30 |
| Fixed (GCE) | 15 |
| Wt NSs (anti-NSs Alexa Fluor 488) | 17 |
| Infected cell (anti-NSs Alexa Fluor 488) | 32 |

**Figure 2.**
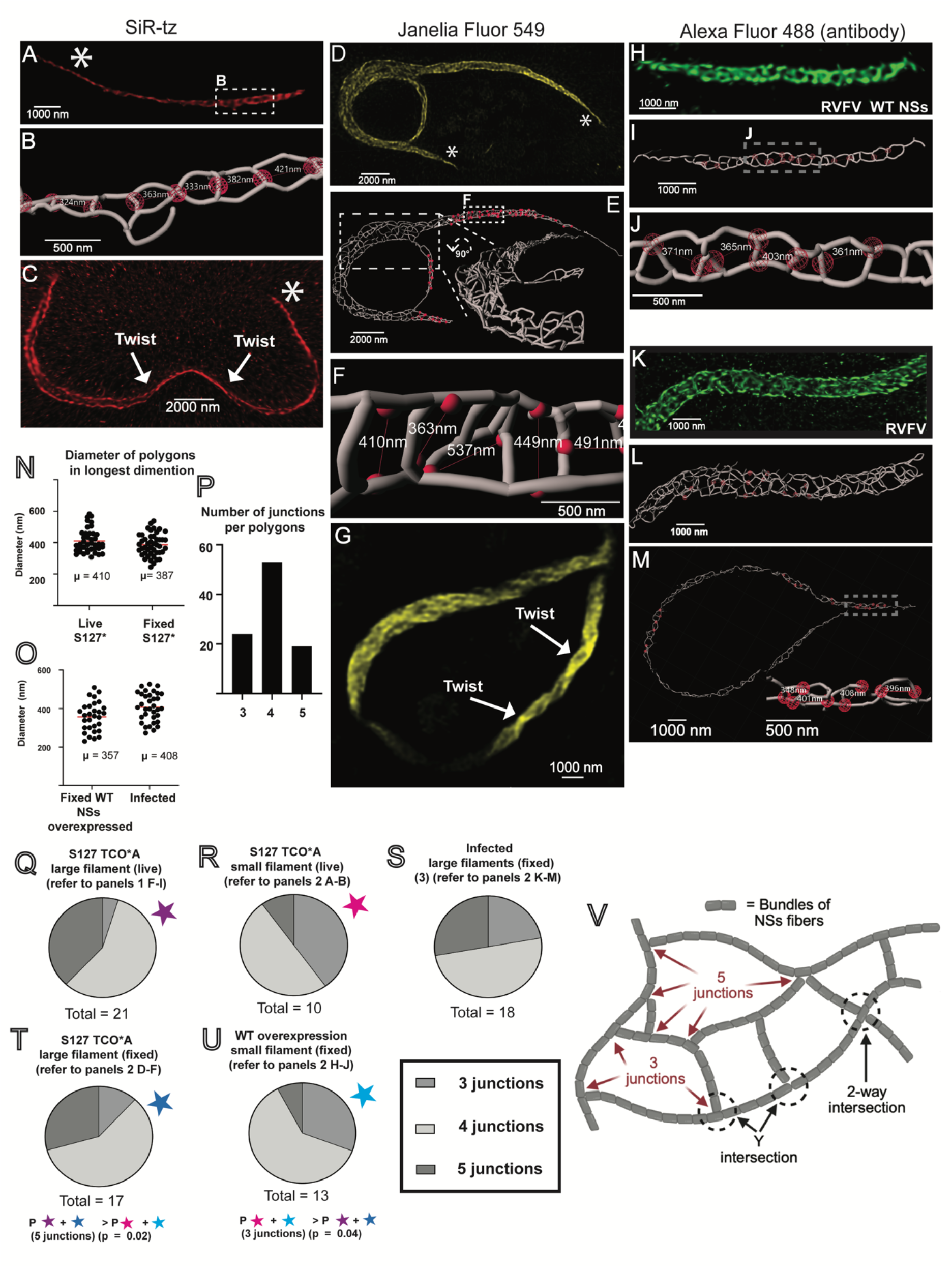
(A) Live-cell 3D-SIM of a small S127* SiR-tz NSs filament in HeLa cells (B) Segmentation with enlarged view of the white dashed inset in A with selected measurement points represented as red spheres. (C) Live-cell 3D-SIM analysis of a small S127* filament in HeLa cells. (D) 3D-SIM analysis in fixed HeLa cells labeled with Janelia Fluor^®^ 549 (yellow). (E) A segmentation of the filament in D. Inset displays the area highlighted within the white dashed box in a more useful perspective and highlights the complex 3D architecture of this filament. (F) Enlarged view of the white dashed inset in E with diameters of the polygons measured in the longest dimension. (G) 3D-SIM of a large S127* NSs filament in fixed HeLa cells. (H) 3D-SIM analysis in fixed HeLa cells of transiently overexpressed WT filaments. (I) A segmentation of H. (J) Enlarged view of the gray dashed inset in I with diameters of the polygons measured in the longest dimension. (K) 3D-SIM analysis of RVFV infected HeLa cells fixed in BSL-3. (L) Segmentation only with selected measurement points represented as red spheres. (M) Segmentation only of a second NSs filament from a RVFV infected HeLa cell, with selected measurement points represented as red spheres. The inset displays the area highlighted in the white dashed box in a more useful perspective. (N) Scatter plot showing diameters of the polygons measured in the longest dimension filaments captured using live-cell (2 filaments) and fixed (1 filament) 3D-SIM. (O) Scatter plot showing diameters of the polygons measured in the longest dimension of filaments imaged in RVFV infected cells (3 filaments) and transiently overexpressed NSs WT fixed (2 filament) 3D-SIM. (P) Histogram showing the number of polygons made up of 3, 4 and 5 junctions. (Q-U) Comparison between number of junctions and overall filament size. Infected fixed large refers to measurements from three filaments imaged in infected cells, of which two are displayed in Figure 2 K-M. (V) Our proposed model for the NSs polygon web architecture.

We defined “sheets” as containing rows of polygons along the long axis of the filament, sometimes in an alternating series **(Figure 1F-I and movie 3)**. In contrast, “chains”, have a single linear row of organised polygons **(Figure 2A-C)**. In addition, we noticed the “sheet” and “chain” morphology would usually collapse at the extreme ends of the fibre into a compressed coil **(Figure 2A, C and D, asterisks, movie 2)**. Importantly, we observed similar architectures in WT NSs filaments in fixed cells. For example, polygon chains were observed in transiently overexpressed WT filaments **(Figure 2H-J)**, and we also observed large heterogeneous polygon sheet and chain arrangements in RVFV infected cells that were deactivated under BSL-3 conditions **(Figure 2K-M) (# of filaments imaged in Table 1)**. The presence of these web architectures in WT filaments confirms these structures are physiologically relevant and not a property of the uAA engineering.

Quantitative analysis, of NSs polygons demonstrated a mean diameter of 0.410 μm and 0.387 μm for **S127TCO*A** live and fixed filaments, respectively, which are consistent with the observed diameters for WT NSs filaments in fixed cells **(Figure2 N-O and movie 3)**. Interestingly, analysis of polygons in all types of filaments revealed that most polygons contained four Y-shaped junctions, with three and five Y-junction-polygons also present **(Figure 2P)**. We wondered whether the morphological differences between small and large filaments were associated with a change in polygon type. In smaller filaments, whether live or fixed, there were 3-4-fold more 3-Y polygons compared to 5-Y polygons. Conversely, larger filaments contained 3-4 fold more 5-Y polygons than 3-Y polygons **(Figure 2 Q-U)**.

## Discussion

Our structural analysis of near-native NSs filaments in live-cells shows an intricate network of NSs fibres that exist with discrete morphologies. Our data suggest that regions where filaments form networks of polygons with higher numbers of junctions offer increased potential for complex network shapes and angles, giving rise to the complex internal structures of larger NSs filaments. Furthermore, given the structural constraints that prevent 70–90° perpendicular Y-shaped branching within continuous filaments, we suggest a model whereby NSs filaments form polygonal webs via Y-shaped intersections, as seen for actin networks^14^ **(see our proposed model in Figure 2V)**. Interestingly, Li et al, captured possible examples of Y-shaped intersections of individual NSs fibres in their recent manuscript (note the criss-crossing NSs intersections in **fig. S1F** ^4^.

It is interesting to speculate small filaments might represent early infection structures optimised for initial host targeting and large filaments could represent mature structures with enhanced capacity for complex regulatory functions. Finally, this architecture suggests RVFV has evolved a sophisticated strategy to compartmentalize the nucleus into functionally distinct zones. Our findings raise important questions about the potential advantages of discrete NSs structural domains for organizing E3 ligase activity: Do different polygon types recruit distinct co-factors? Could this architecture enable localized control of substrate degradation?

## Acknowledgments

We thank David Bhella (CVR) for insightful comments and Iain Harley (CVR) for help with our model presented in Figure 2. We thank Colin Loney and Deirdre McLachlan in the CVR bioimaging facility for their helpful guidance and suggestions. We thank the Beatson Advanced Imaging Resource (BAIR; RRID: SCR_023875). This work was supported by a Future Leaders Fellowship (MR/W010690/1), and the Royal Society Research Grant (RGS\R1\231494) (to S.D.C). LMC is supported by core grant (A17196) to the Cancer Research UK Scotland Institute (Research Organization Registry 03pv69j64) and PT core grant (A31287) to its Core Services and Advanced Technologies (A31287).

## Author contributions

Conceptualization: SDC, BGD, JD; investigation: JD, PAT, SDC; methodology: JD, SDC, PAT, LMC; visualization: JD, SDC; formal analysis: JD, SDC; writing (original draft): JD, BGD, SDC; writing (review and editing): JD, SDC, PAT, LMC.

The authors declare no competing interests.

## Materials and Methods

### Cell lines

HEK-293T (ATCC, CRL-11268), VeroE6, and HeLa cells (ATCC, CCL-2) were maintained in high-glucose DMEM media supplemented with GlutaMAX® and sodium pyruvate (Cat.No.31966021), 10% fetal calf serum (FCS), and 100 U/ml penicillin/streptomycin (p/s). BSRT-7/5 CL21 cells are a clone derived from BSRT-7/5 cells sourced from K.-K. Conzelmann, Ludwig-Maximilians-Universität München, Germany. BSRT-7/5 CL21 and BHK-21 cells were maintained in GMEM (Cat.No.11710035), supplemented with 10% fetal calf serum (FCS), 10% tryptose broth (TPB), and p/s. Additionally, BSRT-7/5 CL21 cells were supplemented with 0.25 mg/ml G418. To minimize background signal, FluoroBrite DMEM (Cat.No.A1896701), supplemented with 10% FCS was used for 3D-SIM imaging experiments. All cell lines in our experiments were cultured at 37°C with 5% CO2.

### Construction of RVFV antigenomic cDNAs

Virus antigenomic cDNAs for RVFV (ZH548 strain) genomic segments (L, M and S) were synthesized with a plasmid backbone that contained constitutive elements, including a colE1 origin of replication and a kanamycin resistance marker, sequences for a T7 promoter, T7 terminator and a hepatitis delta ribozyme (GeneArt, TFS). The sequences of the antigenomic virus segments were obtained from NCBI using the following accession numbers: ZH548 L – NC_014397.1, ZH548 M-NC_014396.1, ZH548 S – NC_014395.1 15. cDNA sequences are available on request.

### RVFV rescue

BSRT-7/5 CL21 cells were seeded in 6-well plates at a density of 7 × 105 cells per well on the day before transfection. Three plasmids encoding the 3 viral segment cDNAs (0.5 μg per plasmid) were transfected using Trans-IT LT-1 (Mirus Bio). On day 3 and day 6, development of visible and highly visible CPE, respectively, was observed and the supernatant was harvested. To confirm the presence of virus and to determine viral titer, a plaque assay was performed using BHK-21 cells. Virus was also passaged in BHK-21 cells using a multiplicity of infection of 0.01 to produce a working stock.

### Tetrazine dyes

Janelia Fluor® 549 was dissolved in DMSO to a concentration of 5 μM and aliquoted for storage at −20°C. For labeling, the stock was further diluted in PBS to a final concentration of 50 nM. SiR-tetrazine (SiR-tz) was dissolved in DMSO to a final concentration of 500 mM and aliquoted for storage at −20°C. For labeling, the SiR-tz stock was then diluted in PBS or cell media to a final concentration of 1 mM. The methyl tetrazine dye CF®500 was dissolved in DMSO to a concentration of 10 mg/ml and aliquoted for storage at −20°C. For labeling, this stock was diluted to 2.5 μg/ml with media before use.

### Antibodies

A polyclonal anti-NSs Ab generated to the ZH548 NSs protein was commissioned from Genscript. A fluorescently labeled version of this Ab (Alexa Fluor®488) was also purchased from Genscript. Secondary fluorescently labeled Abs were purchased from Licor (IRDye® 680RD goat anti-mouse IgG, Cat.No. 926-68070 and IRDye® 800CW goat anti-rabbit IgG, Cat.No. 926-32211). Primary Ab, anti-alpha-tubulin, was used as a loading control (Sigma, Cat.No. T5168).

### Immunofluorescence of NSs in RVFV infected HeLa cells

VeroE6 cells were grown in 8-well chamber slides and infected with RVFV (ZH548) at a multiplicity of infection (moi) of 0.1. After 24 hrs of infection, cells were fixed for 1 hr with 4% formaldehyde and washed twice in PBS before being permeabilized 3 times with 0.1% triton X-100 in PBS on a shaker. The fixed cells were then blocked with 0.5% BSA in PBS for 30 mins at room temperature on a shaker. Following blocking, cells were incubated for 1 hr on a shaker at room temperature with an anti-NSs Alexa Fluor® 488 linked Ab diluted in block solution and washed 3 times for 5 minutes in PBS before immediate analysis of fluorescent staining.

### Genetic-code expansion plasmids

pcDNA3.1(+)_U6 tRNAPyl_CMV NESPylRS(AF) was a gift from Ivana Nikić-Spiegel (Addgene plasmid # 182287; http://n2t.net/addgene:182287; RRID:Addgene_182287). Constitutive over-expression vectors containing the DNA coding sequence of NSs derived from the RVFV strain ZH548 (NC_014395.1, Bird et al., 2007) with amber stop-codon (TAG) substitutions (S99, K108, S112, Q138, Q140, Q199, K202, S206, S226) were purchased from Twist Bioscience (pTwist CMV wPRE BG Neo NSs). A plasmid containing the WT NSs sequence was created using the QuiKChange® protocol (Stratagene). Additional amber stop-codon substitution mutations (Q12, K84, S119, S126, S127 and K150) were also produced in this plasmid.

### Transfection for genetic-code expansion

For wide-field imaging, HeLa cells were seeded at a density of 2 × 10^4^ cells per well in 8-well chamber slides with a glass bottom (Ibidi, Cat.No.80807) or a tissue culture treated polymer surface (Ibidi, Cat.No.80806). For 3D-SIM, glass-bottomed 8-well chamber slides were coated with human fibronectin at 20 μg/ml. Human fibronectin was purchased from Merck (Cat.No. FC010). Slide mounting medium was purchased from Ibidi (Cat.No. 50001). For Western blotting, HEK-293T and HeLa cells were seeded at 4 × 10^4^ cells per well in 24-well plates. Transfection of HEK-293T and HeLa cells in 24-well plates and 8-well chamber slides was carried out using lipofectamine®3000 according to manufacturer’s instructions for 24 hours (TFS, Cat.No.L3000001). Trans-Cyclooct-2-en-L-Lysine (TCOA) was purchased from Sirius Fine Chemicals (SiChem) (Cat.No. SC-8008). A 100 mM stock solution (stored at −20°C) of TCOA was generated by dissolving solids in 0.2 M NaOH and 15% v/v DMSO. Culture media was replaced with media supplemented with TCOA at 250 μM 10 minutes post-transfection.

### Western blotting

RVFV NSs samples for polyacrylamide gel electrophoresis (PAGE) were produced in HEK-293T and HeLa cells. Typically, cells grown in 24-well plates were lysed 24 hr post-transfection using 100–200 μl per well of reducing protein loading buffer before incubation at 95°C for 10 min. Samples were then stored at −20°C before use. Before PAGE, samples were incubated at 85°C for 5 mins prior to gel loading on NuPAGE Bis-Tris gels (Thermo Fisher Scientific, Cat.No NW04122BOX). After approximately 1 hr, the PAGE was stopped and proteins transferred to a nitrocellulose membrane using a semi-dry system with a voltage of 15 V for 30 mins [BioRad Trans-dry system® (discontinued)]. The membrane was stained using a Licor protocol (Intercept®PBS variation) and protein visualized on a Licor Odyssey CLX Imager according to manufacturer’s instructions. Bolt® LDS sample buffer and reducing agent was purchased from Thermo Fisher Scientific (Cat.No. B0007 and Cat.No. B0009, respectively). Chameleon duo pre-stained molecular weight markers were purchased from Licor (Cat.No.928-60000).

### Fixed cells for stain and imaging

Cells were transfected with pTwist CMV wPRE BG Neo NSs and pcDNA3.1 (+)4x (U6 tRNA M15)CMV NESPylRS(AF) in 8-well chamber slides as detailed above in the presence of TCOA and incubated for 24 hr. To wash out the TCOA and reduce background, cell media was removed, and cells were washed twice with pre-warmed media before incubation for 2–3 hrs. Next, the cells were washed twice in PBS before fixation for 20 mins with 4% formaldehyde. The cells were then washed twice in PBS and permeabilized with 0.1% triton X-100 in PBS for 5 minutes, 3 times. Cells were then incubated for 30 mins at 37°C with a tetrazine dye. To wash out unbound dye, the dye solution was aspirated, and cells were rinsed three times in PBS prior to incubation with PBS on a shaker at room temperature for 30 mins. Fluorescence was visualized immediately, or alternatively mounting medium was added, and chamber slides were stored at 4°C in the dark.

### Live-cells 3D-SIM

HeLa cells were transfected with pTwist CMV wPRE BG Neo NSs and pcDNA3.1 (+)4x (U6 tRNA M15)CMV NESPylRS(AF) for 24 hrs as above in the presence of TCOA. To wash out the TCOA, cell media was removed, and cells were washed twice with pre-warmed media before incubation for 2–3 hrs. The media was then replaced with media containing SiR-tz at 1 mM and incubated for 30 mins. For imaging, cells were washed with media and incubated as above for a further 2–3 hrs to remove unbound dye. The media was then replaced with DMEM Fluorobrite containing 10% FCS.

### EPI-fluorescence

To analyze amber-stop-codon suppression by screening for NSs filament formation, chamber slides containing cells expressing RVFV NSs filaments labeled with tetrazine dyes or stained with an anti-NSs Alexa Fluor Ab were analyzed on a Leica DMi8 wide-field microscope.

### 3D-SIM

Samples in chamber slides were imaged using a Zeiss Elyra 7 lattice SIM microscope using Zen Black 3.0 software, with either a 63× 1.4NA oil immersion or 63× 1.2NA water immersion objective. Lattices used were: for Alexa Fluor 488-stained samples G5 (27.5 μm); Janelia Fluor® 549 samples G4 (32 μm) or G5 (27.5 μm if phase modulation was judged to be sufficiently good); SiR-Tz (647 nm) G3 (36.5 μm), all using 13 phases. Z-stacks were captured at intervals of 91–125 nm depending on the excitation wavelength; or for some samples at 55 nm (over-sampling). Lasers (488 nm, 561 nm and 642 nm, all 500 mW) were typically used at 2.5–4% power. Processing was by SIM2 using “standard live” settings of input SNR medium, iterations 16, regularization 0.065, input and output sampling set to ×4, median filter. Grating period was 718.33 nm and resultant xy scaling 0.031 μm. Additional filaments labeled with Janelia Fluor® 549 were imaged on a ZEISS Lattice SIM 3 using Zen Blue software with a 40×/1.4 NA oil immersion objective. Images were captured in SIM-apotome mode, with a 26.8 μm grating and 5 phases. Z-stacks were captured at intervals of 138 nm and a 561 nm laser was used at 5–8% power. Processing was performed using SIM2 in 3D mode with Wiener filter order recombination, a sharpness filter of 10.5746, no regularization, 8 iterations, with ×4 processing and ×4 output sampling.

### AI powered filament tracing using Imaris

Processed 3D-SIM image data files were imported into the Imaris Image analysis programme (version 10.2.0) by converting Zeiss CZI files to Imaris IMS files using the Imaris file converter (version 10.2.0, Oxford Instruments). To reduce background noise and improve image quality before filament tracing, deconvolution was performed in Imaris using robust deconvolution with 20 iterations. The acquisition parameters inputted were TIRF for imaging modality, 1.4 for objective lens numerical aperture, 1.37 for refractive index with oil immersion medium, and 500 nm imaging distance from the coverslip.

The AI powered filament tracer programme in the Imaris software was applied to the 3D-SIM data to automatically detect and segment RVFV NSs 3D-SIM fluorescence. We used the algorithm setting autopath no loops without soma or spines for filament detection. We applied 3D cropping of the region of interest to reduce processing time and selected a microfilament thinnest diameter of 50 nm (size based on filament diameter in 3D-SIM) and omitted the largest filament diameter value. To define NSs fibers, seed points were added manually, and an iterative machine-learning process of adding and removing seed points to fluorescence lines until a converged isosurface was produced and refined enough that further interaction of the tracing algorithm didn’t significantly change its shape. Manual validation was performed by 2 independent observers and all analyses were performed in duplicate. Lastly, for final refinement of the isosurface, we used the fluorescence intensity thresholding tool to allow for manual analysis of 3D-SIM fluorescence. Disconnected segments were removed and individual micro-filaments were added manually.

### Measurements of NSs polygons

We defined a small filament as one composed of one or two rows of polygons along the long axis of the filament. We term filaments with complex heterogeneous architecture as “large”. Measurements of the longest dimension were performed using Imaris. Measurement points were represented as red spheres and positioned in the center of filament isosurface lines that corresponded to the lines of maximal fluorescence intensity. Occasionally, the filament lines did not coincide with fluorescence intensity lines and measurement points were added to the center of the fluorescence intensity lines only, which was a selection option in the filament tracer programme. Filament isosurface and fluorescence intensity line overlays displayed using the Imaris software were also used to count the number of junctions around hole shapes for each filament labeling type.

